# The *Disc1* deletion common to many inbred mouse strains has negligible effects on social behavior of 129S4 mice in a semi-natural environment

**DOI:** 10.1101/2025.03.17.643700

**Authors:** Olivia Le Moëne, Max Larsson, Walker S. Jackson

## Abstract

Numerous mouse models have been engineered to carry alterations to the Disrupted in Schizophrenia 1 (*Disc1*) gene, thought to be involved in neurodevelopmental conditions. However, most Swiss and probably all 129 mouse substrains, which are widely used in biological research, naturally carry a 25 bp deletion in *Disc1* exon 6. Despite the prevalence of these strains, little has been done to characterize the extent to which this natural mutation affects behavioral output and may unintentionally impact studies. Here, we report on experiments to test the effects of this deletion on social and exploratory behaviors. To model natural conditions, we designed a seminatural environment to house groups of mice (4 females and 1 male; 3 groups per strain) for prolonged periods and then employed this model to study social behaviors. First, we compared behavioral phenotypes in C57Bl/6Jrj (B6) mice and 129S4 (S4^Disc1−/−^) natural mutants to validate our setup. Then, to assess the contribution of the naturally mutated *Disc1* to social behavior differences, the wild-type (WT) *Disc1* allele was crossed into S4 mice (S4^Disc1+/+^). S4 and B6 lines were drastically different, with S4 mice being hypoactive, less explorative, and less social than B6 mice. However, S4 mice expressing WT *Disc1* only marginally differed from S4^Disc1−/−^ mice, showing little to no contribution of *Disc1* to their behavioral phenotype. Thus, this mutation holds little significance for natural exploratory and social behaviors in the seminatural environment.

## Introduction

Millions of people worldwide are affected by neurodevelopmental conditions such as bipolar disorder, major depression, or schizophrenia. Some of these conditions are partly explained by genetic causes. In particular, a Scottish family suffering from a high proportion of psychiatric disorders was discovered to carry a gene translocation (1q42; 11q14.3) on chromosome 1 (St Clair et al., 1990). This gene, later called Disrupted in Schizophrenia 1 (*DISC1*) codes for a protein involved in a variety of neural processes (Brandon et al., 2009), and mutations cause increased susceptibility to psychiatric disorders (Facal and Costas, 2019). The most common *DISC1* mutation in humans is a consequence of a balanced chromosomal translocation (1q42; 11q14.3) that disrupts *DISC1* at intron 8 (Blackwood et al., 2001).

Genetic engineering of mouse models has helped reveal the functional role of a plethora of genes in neurodevelopmental conditions, from ontology to behavioral output. There are at least three different *Disc1* mutations currently in use in mice in which different exons in the endogenous *Disc1* gene were modified (Koike et al., 2006;Kuroda et al., 2011;Shahani et al., 2015). Mouse studies based on these different mutants were intended to map the effect of *Disc1* partial or complete deletion, to establish its role in cognition, emotion and sociality, aspects of personality that are strongly affected in schizophrenia disorders. Indeed, such mutants show multiple alterations in brain development including gross brain morphology, protein expression, synaptic signaling, and brain plasticity (Greenhill et al., 2015;Juan et al., 2014;Koike et al., 2006;Tomoda et al., 2017). Behaviorally, *Disc1* mutants have been reported to exhibit a variety of abnormalities in cognitive, locomotor and sensory abilities, as well as in social, sexual or anxiety-related behaviors. *Disc1* mutants were reported to have deficits in working memory (Juan et al., 2014;Koike et al., 2006), and several strains showed hypoactivity and social impairments (Clapcote et al., 2007;Shen et al., 2008). Effects on spatial memory, motivational and locomotor behavior are sparser and strongly rely on the exact length and location of the mutation (Clapcote et al., 2007;Juan et al., 2014;Koike et al., 2006;Shen et al., 2008).

Buried within this general knowledge is the complication that the functional impact of one of the most pervasive mutations is incompletely understood. Specifically, certain strains of mice naturally carry a 25 bp deletion in *Disc1* exon 6, and the resulting frameshift leads to a stop codon in exon 7 (Koike et al., 2006). Transcripts carrying this deletion should encode a Disc1 protein fragment consisting of the first 528 amino acids of the normal 852 amino acids, plus 13 new amino acids. A first impression conjures the notion that this mutation is like that found in the Scottish family in which the *DISC1* mutation was initially identified and that presented a high number of individuals with mental illnesses. However, the mouse *Disc1* gene encodes multiple mRNA splice variants and the exon 6 with the deletion can be excluded by alternative splicing, resulting in a Disc1 protein variant that encompasses most of the full-length protein (Ishizuka et al., 2007). Importantly, the Disc1 protein in brains of mice carrying this *Disc1* deletion appears to be expressed at relatively high levels and can be bound by many of the same antibodies that bind to the C-terminus of wild-type Disc1 (Ishizuka et al., 2007). Therefore, this mutation is quite unlike the Scottish mutation where none of the C-terminus is expressed, and it is not clear to what extent the mouse *Disc1* deletion has impaired Disc1 function. Nonetheless, this natural deletion has been implicated in causing brain morphology, neuronal development, and behavior abnormalities in mice (Baskaran et al., 2020;Holley et al., 2013;Juan et al., 2014). There are other discrepancies too. Mice homozygous for the natural *Disc1* deletion were indistinguishable from mice homozygous for wild-type *Disc1* when comparing sleep patterns and responses to sleep deprivation (using the gentle handling method for six hours) as measured by electroencephalography and electromyography (Dittrich et al., 2017). However, it was reported that mice heterozygous for this deletion responded to sleep deprivation (using the multiple platform method for 72 hours) with increased microgliosis and reduced induction of brain-derived neurotrophic factor (Tsao et al., 2022). Since many inbred strains carry the natural *Disc1* deletion, including most Swiss-derived and probably all 129 substrains, and many genetically modified mice carry complete or partial genome sequences from these strains, there is concern that some genetically engineered mouse lines may carry *Disc1* deletions unknown to the experimentalists (Clapcote and Roder, 2006). Therefore, it is crucial to determine if the natural *Disc1* deletion may unintentionally affect neurobiological studies. Because of this, and the connection of *Disc1* to neuropsychiatric problems, we sought to understand if this deletion broadly affects social behaviors.

To test for the effect of the natural *Disc1* deletion on social behaviors in a context modeling natural conditions, we established behavioral phenotypes of C57Bl/6Jrj (B6) and S4 mice in a seminatural environment (SNE). This step established that our SNE setup can detect social differences between groups of mice. We then evaluated S4 mice in which the natural *Disc1* deletion allele was replaced with the wild-type (WT) *Disc1* allele, to assess the contribution of the naturally mutated *Disc1* in the initially established behavioral differences. Based on previous findings, we hypothesized that B6 mice would display more exploratory and more anxiety-related behaviors than S4 mice, and that the latter would exhibit altered social behavior. In addition, we expected that if the *Disc1* deletion causes social behavior differences in S4 mice, replacing it with a wild-type version would diminish them.

This study showed that when housed for a prolonged time in a semi-natural environment, B6 mice were surprisingly active, both during the light phase, when nocturnal animals tend to rest, and in the open field, which mice tend to avoid. In contrast, S4 mice were far less active than B6 mice, including reduced exploration and social interactions, and avoidance of the open field. Replacing the mutant *Disc1* gene with a wild-type *Disc1* allele did not change the overall phenotypic picture of S4 mice, including social interactions.

## 1. Material and Methods

### 1.1 Subjects

The mouse lines used in the present experiment were wild-type C57Bl/6Jrj (Janvier laboratories, France, hereafter B6), as well as a commonly used 129 substrain, 129S4 (formerly known as 129/SvJae, hereafter S4 or S4^Disc1−/−^) bred in house. In addition, S4 mice congenic for the wild-type *Disc1* allele (S4^Disc1+/+^) were bred in house, following a procedure described earlier (Dittrich et al., 2017). Briefly, a wild-type *Disc1* gene was introduced into the S4 genetic background by mating S4 mice with C57Bl/6NTac mice, followed by backcrossing to S4 mice. At least five consecutive generations of father-to-son transmission ensured the X chromosomes and mitochondrial genomes were S4 derived, and a transmission from mother to son ensured the Y chromosome was S4 derived. A SNP (single nucleotide polymorphism) analysis at generation 8 revealed that 345 of 347 (99.65%) discriminating SNPs were homozygous for the S4 allele, and the other two SNPs were heterozygous (in *Disc1* heterozygotes) and closely linked to *Disc1.* The line was further backcrossed for 4 additional generations.

The study included a total of 45 mice (36 females and 9 males, 20.9 ± 0.3 and 26.1 ± 0.9 g upon the beginning of the experiment, respectively). The mean age at the beginning of the experiment was 8.7 ± 0.1 weeks for S4^Disc1−/−^, 9.7 ± 0.1 weeks for S4^Disc1+/+^ and 10.2 ± 0.1 weeks for B6. Animals were housed in same-sex groups of 2 to 4 animals in standard cages (mouse 500, NexGen, Allentown Inc., PA, USA) containing wood chip bedding and enriched with nesting material, wooden biting sticks and paper rolls. The lab conditions offered a 12D:12L light cycle, lights being on from 7:00 to 19:00, with an average temperature of 22 °C and a stable humidity of 55%. Food and water were available ad libitum to the mice.

All the experiments performed in this study were approved by Linköping’s Animal Care and Use Committee (Linköpings djurförsöksetiska 7810-2021), and in agreement with the European Union council directive 2010/63/EU.

### 1.2 Apparatus

The seminatural environment (SNE) consisted of an open field (H50 x L45 x W30 cm) communicating with a burrow area through two small doors. The burrow area was composed of a main nest box (H10 x L15 x W15 cm) linked to three small ones (H10 x L8 x W8 cm) by circular corridors of 4 cm diameter (GEHR, PA, USA). Both the open and burrow areas were made of opaque black Plexiglas, while the corridors and nest boxes were made of transparent one (Plexigruppen, Piteå, Sweden). Enrichment was provided in the form of nesting material, 4 wood biting sticks, 5 mats of non-woven fibers, and 2 red polycarbonate rodent tunnels. The same light cycle, humidity and temperature as previously mentioned applied. The burrow area was maintained in complete darkness with the help of an infrared transparent black lid (962 perspex IR), while the open field was not covered (Fig.1A-B).

**Figure 1.**
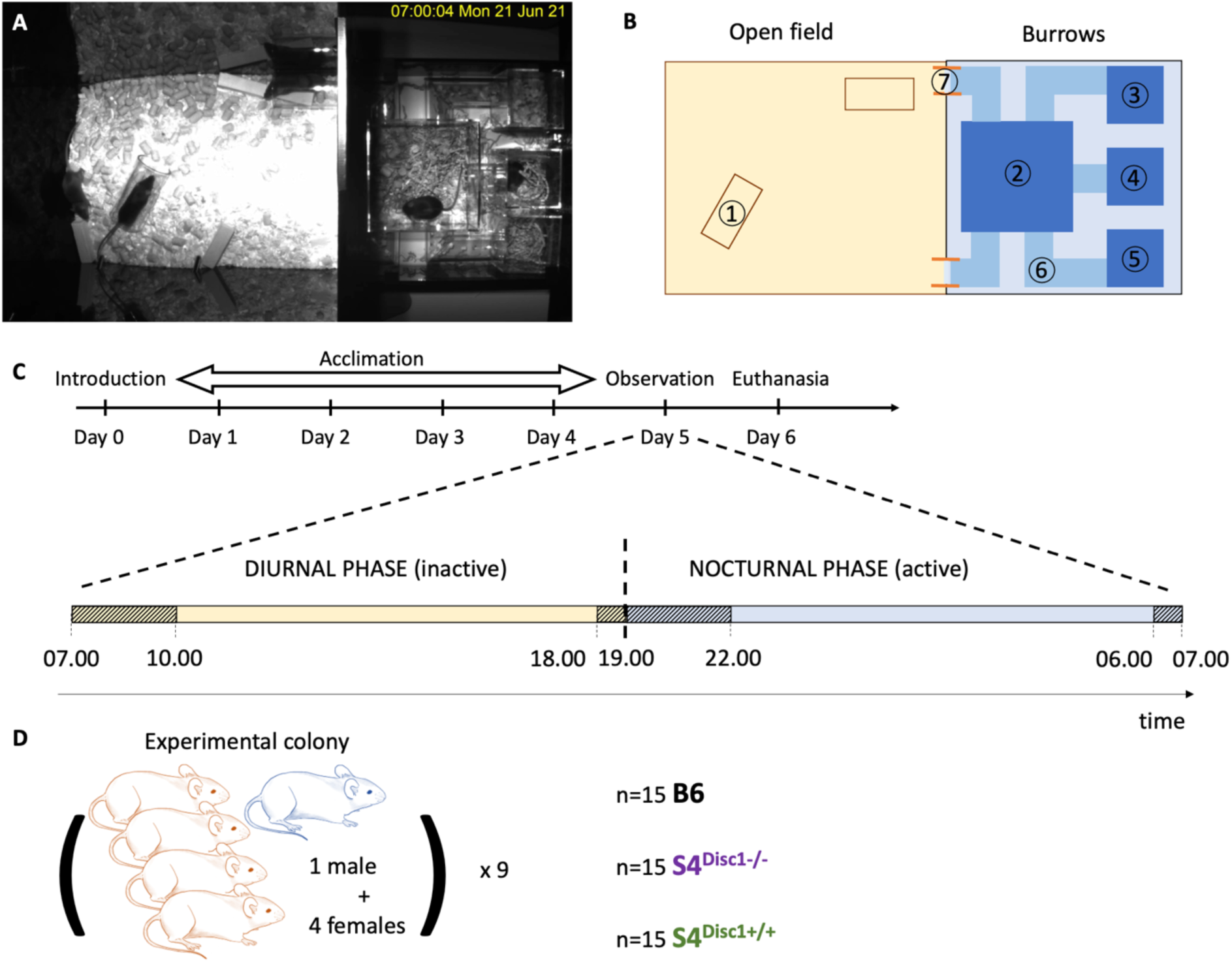
Experimental design. A. Picture of the seminatural environment (SNE) as recorded from above. B. Division of the SNE in zones. Open field: 1=tunnel. Burrows: 2=main nest; 3=sub-nest 1; 4=sub-nest 2; 5=sub-nest 3; 6=corridors; 7=opening. C. Distribution of the observation time during the 12L:12D light cycle on day 5 of the experiment. Observation time is visualized by the striped pattern. Behavior was scored for 4 hours during the diurnal phase and 4 hours during the nocturnal phase. D. Composition of experimental colonies.

In addition to the light cycle, an infrared lamp provided sufficient light for video recording in both areas. A Basler camera (ace 2 USB 3.0) filmed the entire SNE from above, and video recordings were stored on an external hard drive using the Open Broadcaster Software (https://obsproject.com). [FIGURE1]

### 1.3 Procedure

Prior to introducing each experimental colony of mice to the SNE, its floor was covered with a thick layer of wood chips bedding and provided with above-mentioned enrichments. Two spouts in the wall connected to bottles of 0.5 L of water and approximately 1 kg of food pellets were available in a corner of the open field of the SNE. Males and females were mixed as to obtain a more representative observation of natural social interactions. Most often, all 4 females came from the same home cage, while a single male was chosen from another cage. We believe that after 5 days in the SNE, all animals were familiar to each other and the group stabilized, as highlighted by other studies conducted in the Visible Burrow System (VBS) (Bove et al., 2018) as well as an in-house pilot study (unpublished) that showed stabilization of behavior and social interactions by day 3. After each experimental session, the SNE was emptied and disinfected before starting a new colony.

On day 0 at 10:00, the mice were briefly anesthetized with isoflurane (< 1 min/animal) and shaven on the back with different patterns for identification purposes on video recordings. They were then released into the SNE. The experiment was terminated on day 6 at 10:00, when the mice were sacrificed using carbon dioxide asphyxiation followed by cervical dislocation.

### 1.4 Design

Each colony established in the SNE consisted of 4 females and 1 male. The 4 females came from the same cage and thus were familiar to each other. The colony was left undisturbed for the whole duration of the experiment (Fig1.D).

### 1.5 Behavioral observations

In order to minimize the observation of novelty- and socially-induced stress behavior, we conducted behavioral observations on day 5, after 5 full days of acclimation. This is considered enough for the colony to be established and the mice familiar with the environment (Bove et al., 2018). We observed most aggressive behavior in the first hour following release into the SNE, but overall aggression levels were low. Following extensive observation of the video record, we established an ethogram for mouse behavior scoring in the SNE (Table1). The ethogram used here was established on the basis of others used for mice hosted in VBSs and for rats observed in SNEs. We have also conducted other studies on colonies of B6 mice with the current ethogram which are currently under review or unpublished. All data were scored by a single, experienced investigator who was blinded to the genotypes of the S4 strains but not the B6 strain. Behaviors were scored by scan sampling, taking a 1-min sample at 10 min intervals during 8 hours in total, 4 of which were during the dark phase (Fig.1C). Scoring was made manually with the Observer XT 15 software (Noldus, Wageningen, the Netherlands). Locomotor activity was assessed by proxy by counting how often successive behaviors were displayed in a different area of the SNE. Since the zone of the SNE where each behavior was observed was recorded in the coding scheme, each time a behavior was displayed in a different zone than the previous one, this was counted as a zone transition. An in-house study was performed to accurately score the number of demarcation lines crossed by the mice, and a comparison with the “by proxy” method showed a 79% correlation (Pearson’s correlation, p< 0.01, unpublished data).

### 1.6 Statistical analysis

Behavioral occurrences were summed over the entire duration of the observation period, thus 24 min of observation each for the diurnal and the nocturnal phases. The diurnal phase was the sum of 3 hours following dawn (7.00-10.00) and 1-hour preceding dusk (18.00-19.00). Similarly, the nocturnal phase was composed of the 3 hours following dusk (19.00-22.00) and the hour preceding dawn (6.00-7.00). This was done for all the behaviors mentioned in the ethogram but resting behaviors (resting alone, pair resting, huddling), for which durations, but not occurrences, were summed up. This was because the time spent sleeping was more meaningful than the number of occurrences of falling asleep.

We first analyzed the differences between all 3 mouse lines using one-way ANOVAs, or Kruskal-Wallis tests when relevant. Posthoc comparisons were obtained with the Tukey or the Dunn’s test. To confirm these results with an alternative analysis, we also calculated differences between B6 and S4^Disc1−/−^ for each behavior using Student’s *t*-tests or Mann-Whitney U tests when data deviated from the normal distribution. We then conducted the same analysis comparing S4^Disc1−/−^ and S4^Disc1+/+^ mice in which the wild-type Disc1 had been crossed into the S4 background. When the data deviated greatly from the normal distribution according to the Shapiro-Wilk’s test, we analyzed the effect of Disc1 replacement with Wilcoxon tests and compared the mouse lines with the Kruskal-Wallis ANOVAs.

Finally, locomotor activity and weight gained by the end of the experiment were compared among all 3 experimental groups with a one-way ANOVA. Locomotor activity was estimated by the number of zone transitions in the SNE (see zone division in Fig.1B). Statistical analysis and data figures were obtained using GraphPad Prism 9.1.2 and R 4.1.3.

## 2. Results

### 2.1 Behavioral phenotypes of B6 and S4^Disc1−/−^ in the seminatural environment

Since there are thousands of protein coding differences between 129S5 and B6 (Timmermans et al., 2017) and 129S5 is closely genetically related to S4 (Simpson et al., 1997;Threadgill et al., 1997), there are likely thousands of protein coding differences between S4 and B6. Thus, behavioral differences between S4 and B6 are likely controlled by a combination of genetic factors and they may not involve the *Disc1* alleles. We performed this initial comparison between S4 and B6 strains as we predicted there would be differences detectable using our SNE setup. A negative result from this initial comparison would indicate the SNE is not a sensitive tool for measuring social differences, and any negative results for the comparisons between the S4 *Disc1* genotypes would be meaningless.

Consistent with observations made in other contexts (Dittrich et al., 2017;Kaczmarczyk et al., 2021) S4^Disc1−/−^ mice were less active than B6 mice in the seminatural environment, during both the diurnal and nocturnal phases of the light cycle, with reduced exploratory and social activity, and increased resting time (Fig. 2). During the diurnal phase, when mice normally rest, S4^Disc1−/−^ mice displayed less exploratory sniffing and digging, as well as less social approaches, agonistic nose-off episodes, and food disputes than B6 mice. In addition, compared to B6 mice, S4^Disc1−/−^ mice also spent more time resting alone but, interestingly, not huddling nor resting in pairs (Fig2.A).

**Figure 2.**
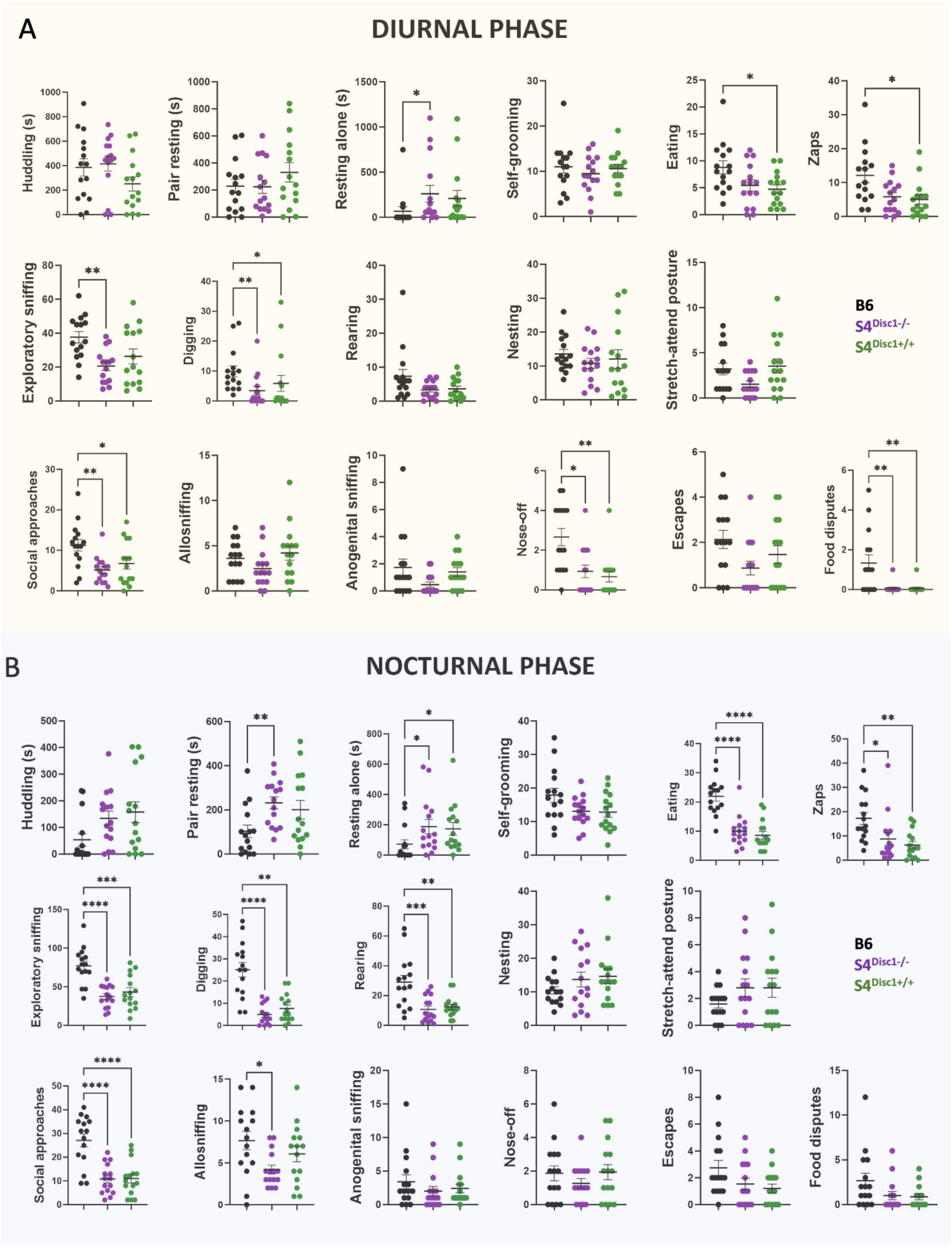
Comparison of maintenance, exploratory and social behaviors in Bl6 mice, natural S4^Disc1−/−^ mutants (n=15) and S4^Disc1+/+^ mice (n=15). Data are behavioral occurrences, except for resting behaviors (huddling, pair resting and resting alone), expressed in duration (s). Data are mean ± SEM. One-way ANOVAs or Kruskal-Wallis tests, * p < 0.05, ** p < 0.01, *** p < 0.001, **** p < 0.0001.

During the active, nocturnal phase of the light cycle, S4^Disc1−/−^ mice routinely expressed less exploratory sniffing, digging, rearing, and eating than B6 mice. They also emitted less pro-social behaviors such as social approach and allosniffing than B6 mice. There was no difference in agonistic behaviors. Sleeping-related behaviors including resting in pairs or resting alone were increased throughout the nocturnal phase in S4^Disc1−/−^ mice (Fig.2B).

Behaviors commonly associated with anxiety, such as self-grooming, nesting, or displaying a stretch-attend posture, did not differ between lines (all ps>0.052), but zaps, single jolting movements that are unprovoked, were less frequent in S4^Disc1−/−^ mice during the nocturnal phase (Fig.2A-B). In summary, our SNE setup can reveal several social behavioral differences between groups of mice.

### 2.2 Disc1 replacement does not modify S4 behavioral patterns

We predicted that if the *Disc1* deletion strongly affects these behaviors in S4 mice we should be able to detect differences when comparing S4^Disc1+/+^ and S4^Disc1−/−^ mice. Surprisingly, S4^Disc1+/+^ mice were nearly identical to S4^Disc1−/−^ mice, regardless of the light cycle phase (Fig.2A-B). The groups were not different in any of the 17 measured behaviors for either light phase. Moreover, S4^Disc1+/+^ differed from B6 in a pattern nearly identical to that for S4^Disc1−/−^ mice. Among the nine behaviors in which S4^Disc1−/−^ mice differed from B6 mice during the nocturnal phase, seven were also changed in the same direction in S4^Disc1+/+^ mice. Similarly, six behaviors were changed for both S4 genotypes during the diurnal phase, four of which were the same. The overall behavioral profiles are summarized in Fig. S1. The time of day when each behavior occurred was similar between all lines (Fig.S2). This analysis revealed differences between the B6 line and both S4 lines, but not between the S4 lines themselves.

The above results suggested there are no strong differences between the S4 lines, but we wondered if removing data from males or applying an alternative statistical method might reveal weak differences. We therefore reanalyzed the data from females only and, despite the reduced statistical power with fewer replicates, that analysis found differences between B6 and the S4 strains that were nearly the same as the first analysis (Table S1). Furthermore, this female-only analysis revealed only a single behavior, huddling, where S4^Disc1−/−^ and S4^Disc1+/+^ mice differed (Table S1). We also reanalyzed the complete data set using Student’s *t*-tests or Mann-Whitney U tests. For the S4^Disc1−/−^ and B6 comparisons nine behaviors were different in the diurnal phase and ten in the nocturnal phase (Fig.S3). In contrast, only huddling during the diurnal phase was different between S4^Disc1−/−^ and S4^Disc1+/+^ mice (Fig.S4). Therefore, S4^Disc1−/−^ and S4^Disc1+/+^ mice do not have strong social behavior differences when measured in the SNE.

### 2.3 Locomotor activity of B6, S4^Disc1−/−^ and S4^Disc1+/+^ in the seminatural environment

Locomotor activity in the SNE was assessed by the number of scored transitions between the zones inside the two main areas of the SNE: the open field (OF) and the burrow system (B). The total number of transitions (OF+B summed) was lower during the diurnal than the nocturnal phase (F_(1, 42)_=10.820, p=0.002), but showed no effect of the mouse line (F_(2, 42)_=1.740, p=0.188) nor any interaction between these factors (F_(2, 42)_=0.948, p=0.396) (Fig.3A). However, when looking at the spatial distribution of the locomotor activity, we found a strong interaction between the mouse lines and the area of the transitions (F_(2, 42)_=90.940, p<0.001). Post hoc tests revealed that B6 mice were more active in the OF than both S4 strains (p<0.001 in both cases). Conversely, B6 mice were less active in the burrows than both S4 strains (p<0.001 in both cases) (Fig.3B). In addition, there was a larger number of transitions in the OF than the B (F_(1, 42)_=45.350, p<0.001), but no main effect of the mouse line (F_(2, 42)_=1.740, p=0.188).

**Figure 3.**
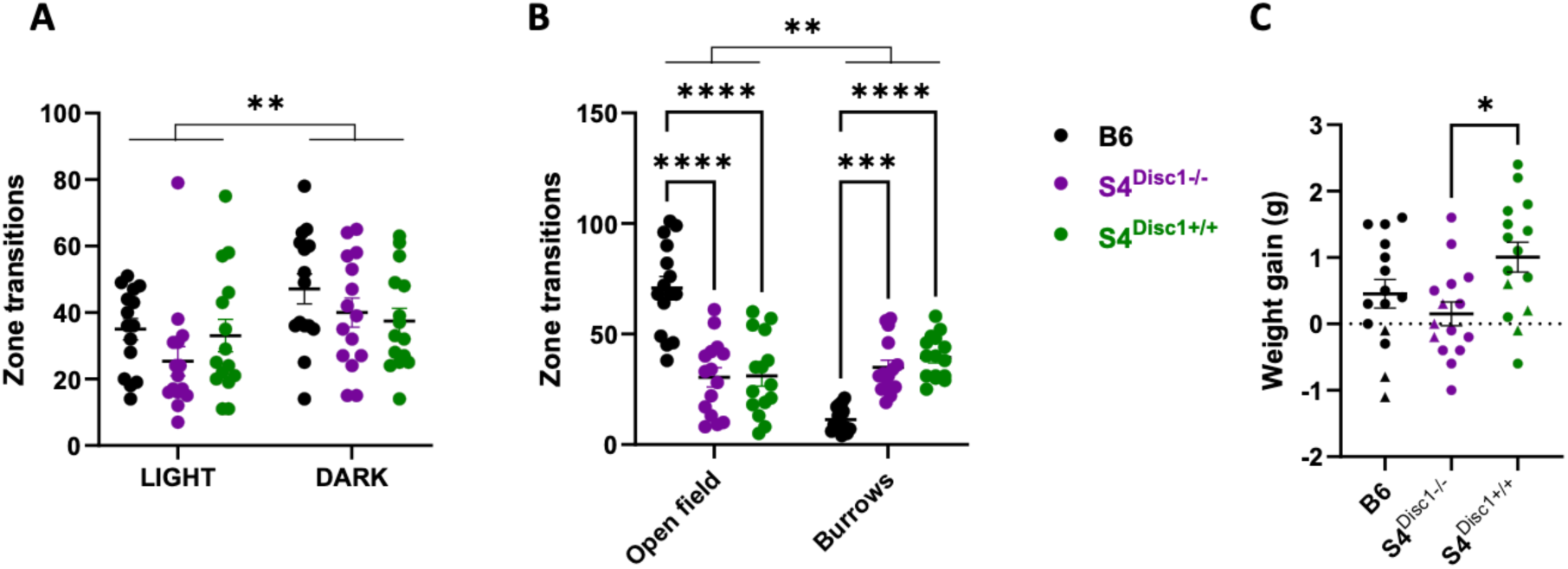
Locomotor activity and weight changes in the SNE. A. Total number of zone transitions (in the open field and the burrow system) over the diurnal (light) and nocturnal (dark) observation period. B. Spatial distribution of zone transitions (diurnal and nocturnal phases summed) between the open field and the burrows. C. Weight gain (g) after 7 days in the SNE. Males are indicated by a triangle symbol. Analysis of female only data changed the significance of the weight gain difference between S4^Disc1−/−^ and S4^Disc1+/+^ mice to be **. Data are mean ± SEM. Two-way ANOVAs with repeated measures (A-B), one-way ANOVA (C). **, p< 0.01 *** p < 0.001, **** p < 0.0001.

Finally, body mass changes can be caused by altered behaviors, among other reasons. By the end of the experiment, weight gain during their 7-days stay in the seminatural environment was significantly more in S4^Disc1+/+^ mice (1.0 g) than in S4^Disc1−/−^ (0.2 g). Despite B6 mice being scored for nocturnal eating a lot more than the S4 groups, the weight gain in B6 (0.5 g) did not differ from the S4 groups (Fig.3C).

## Discussion

This is the first study to characterize S4 behavioral phenotypes in naturalistic conditions. We found that S4^Disc1−/−^ mice differed from B6 in the seminatural environment in many ways. S4^Disc1−/−^ spent more time resting, and less time exploring their environment. They performed fewer occurrences of social investigation and appeared socially withdrawn. Surprisingly, S4^Disc1+/+^ did not differ from S4^Disc1−/−^, except for a recovery in anogenital sniffing to levels comparable to those expressed by B6 mice, only during the diurnal phase, and a significant weight gain. Neither S4^Disc1−/−^ nor S4^Disc1+/+^ mice showed any clear difference in the overall locomotor activity compared to B6. However, both S4 strains showed a preference for activity in the dark burrows compared to the light-exposed open field, opposite to how B6 mice behaved. In hindsight, the lack of difference between S4^Disc1+/+^ and S4^Disc1−/−^ mice could be expected since the deletion mutation only partially disrupts the transcript and much of the protein is still expressed at near normal levels (Ishizuka et al., 2007).

The ability of SNEs, or visible burrow systems, to differentiate strain differences in social behavioral phenotypes is well established (Pobbe et al., 2010) and corroborated in the different behavioral profiles of B6 and S4 mice found in the present study. We therefore considered it valuable to use the same approach to study this single gene mutation. We believe that the results are meaningful in two ways: first, because the individuals can express this mutation-induced difference throughout a larger range of behaviors, which might be undetectable in standard tests. Second, because of this larger behavioral repertoire, individuals can also implement compensatory mechanisms that could mask a potential difference. This is meaningful because any detected difference should therefore be considered biologically significant.

Therefore, the lack of a *Disc1* effect in the present study could result from it being masked by compensatory mechanisms implemented in the SNE and unavailable in simple tests. We would argue that this is crucial, and that if a difference is masked in more ecologically valid procedures, this is highly relevant for result translatability, and clinical observations. Likewise, one or more other genes in S4 could work synergistically with the *Disc1* deletion to cause schizophrenia-like abnormalities. However, if S4 mice are characterized as having a schizophrenia-like phenotype, then it exists independently of the *Disc1* deletion. Thus, the genetic background of S4 mice may mask effects of the *Disc1* deletion.

Though our findings are consistent with previous studies reporting hypoactivity (Shen et al., 2008) and reduced social interactions in some *Disc1* mutants (Clapcote et al., 2007), surprisingly, introducing a WT *Disc1* gene into the S4 strain did not change these features. The *Disc1* gene was initially thought to be a direct risk gene for schizophrenia, but more recently, the strength of this association has been questioned (Farrell et al., 2015;Mathieson et al., 2012). Nonetheless, Disc1 interacts with a great number of proteins and is involved in many neuronal processes. It is now thought that Disc1 acts as a molecular scaffold to regulate molecular and physiological protein expression (Tomoda et al., 2017), and that these actions, in combination with environmental stressors, can lead to the development of psychiatric symptoms categorized as schizophrenia. Thus, mice raised similarly and being hosted in a large, familiar environment with limited stress exposure might not express behavioral alterations the way mice tested in new, anxiety-inducing procedures would.

Why are there seemingly contradictory conclusions about the effects of the natural *Disc1* deletion? Studies by Gogos and colleagues employed a mouse line genetically engineered to model the Scottish deletion (Koike et al., 2006). The manipulation included a stop codon in exon 8 and a transcription terminating sequence in intron 8. Mice carrying this engineered allele were found to have mild changes to cognitive behaviors and neuronal architecture and plasticity (Koike et al., 2006;Kvajo et al., 2008). However, embryonic stem cells isolated from 129S6 mice were used for the manipulation, and thus, the engineered allele also included the natural 25 bp *Disc1* deletion in exon 6 resulting in a premature stop codon in exon 7. Since the deletion was upstream of the engineered translation and transcription stop signals, it was assumed to supersede the engineered modifications (Koike et al., 2006). Soon after, the 25 bp deletion was reported to be in many 129 inbred lines, raising the concern that it might have unexpected consequences on experiments involving gene-targeted mice derived from 129 embryonic stem cells, which is the case for most such mouse lines. The concern was further escalated when the same 25 bp deletion was found in other widely used strains of mice (Ritchie and Clapcote, 2013). However, confusion was added to the story when it was reported that many antibodies that were validated to bind to recombinant mouse Disc 1 protein, including two targeting sequences that are downstream of the 25 bp deletion, resulted in indistinguishable bands on western blots of brain lysates derived from B6 and 129S6 mice (Ishizuka et al., 2007). It was subsequently found that the mice genetically engineered to have the exon 8 stop codon and intron 8 transcription termination signals express only the N-terminal portion of Disc1 (Kvajo et al., 2008). Since mice carrying the unmodified *Disc1* allele with the natural 25 bp deletion appear to express most of the same Disc1 forms as mice carrying the wild-type gene, readers must carefully read the details of research articles to understand which *Disc1* allele is under investigation. Furthermore, since we see strong differences between standard S4 and B6 mice, readers should be aware if comparisons reporting differences due to *Disc1* variants are done in the same genetic background or are instead comparing two different strains of mice. For example, compared to B6 mice, BTBR T+ Itpr3tf/J mice (considered a model of autism) displayed differences in VBS (Bove et al., 2018) and behaviors reminiscent of autism in a group-housed setting (Endo et al., 2019). Although BTBR T+ Itpr3tf/J mice carry the same *Disc1* deletion as S4 mice, those studies did not compare *Disc1* variants in the same genetic background and the authors were careful not to make claims that the *Disc1* deletion was the reason for the differences. In contrast, another study comparing mouse strains with different genetic backgrounds, one of which harbored the *Disc1* deletion, attributed differences to the *Disc1* deletion (Sultana et al., 2018).

Finally, studies using the opposite approach to ours, moving the unmodified *Disc1* deletion into the B6 background, report that the natural *Disc1* deletion affects many neuroanatomical and behavioral phenotypes (Baskaran et al., 2020;Holley et al., 2013;Juan et al., 2014;Tsao et al., 2022). Thus, had we used that combination of B6 mice, it is possible we would have detected a stronger effect of the *Disc1* deletion. We considered also doing that experiment but, given the results we obtained with the S4 comparisons, had there been a difference detected in B6 strains, the conclusion would be that there is not a strong, dominating effect from the mutation. It was therefore difficult to justify the extra mice needed to import and breed a B6 *Disc1* deletion line for that single experiment.

### Implications for S4 mice and B6 - which one should be considered “normal”?

We previously observed variable hyperactivity behaviors including jumping thousands of times or walking thousands of meters, within a 12-hour period in a standard mouse cage, by B6 but not S4 mice (Kaczmarczyk et al., 2021). In the current study, we again found drastically different phenotypes in B6 and S4 mice, which appear to be unrelated to the *Disc1* mutation carried by S4 and many other inbred mouse strains. Considering the widespread use in neuroscientific and biomedical research of mice naturally carrying this mutation, both known and unknown, it is reassuring to find that this mutation should hold no significance regarding natural exploratory and social behaviors. Furthermore, when a mouse strain other than the typical B6 is desired, the S4^Disc1+/+^ mice can be used to avoid concerns about the *Disc1* deletion.

A potential limitation of our study is that the groups comprised individuals of the same strain. It is possible that mixing mouse strains would have revealed other differences, for example, preferred social interactions between the same or different strains, or social buffering of the observed differences. Furthermore, a larger number of replicates may have revealed subtle differences between the S4 strains that our study did not detect. While possible, we aimed to detect strong differences, like those revealed in comparisons of B6 and S4.

Another limitation is that the SNE has not yet been widely used. While we could not conduct a strict comparison of the equivalence between social behaviors observed in standard tests and those observed in the SNE, as those are by nature impossible, several factors suggest that those should be highly correlated. For example, using home cage monitoring systems (HCMS), other research teams have found similar locomotor activity as in standard tests (Tang et al., 2002). Moreover, standard anxiety tests can yield conflicting results, but richer environments can simplify their interpretation (Grieco et al., 2021). In the last two decades, there have been consistent efforts to use home cage monitoring and more naturalistic settings to reduce stress and obtain more ecologically-valid results. The present study contributes to this effort by creating a data continuum from individualized standard test, to home cage monitoring and toward SNE monitoring and testing.

Our study included groups with mixed sexes since this is more natural. Male mice are territorial and would fight, sometimes to death, if they are confined in a space where females are also present. Therefore, it was not ethically possible to include more than one male per group. However, female-male interaction is part of the natural social organisation of mouse groups, therefore it would have been reductive to observe single-sex groups. Comparing female-female and female-male interactions would lead to very imbalanced group size due to the current sex ratio and therefore the results would not be sufficiently robust. Since this experiment was performed on groups of individuals with the same genotype, the conclusion was drawn at the group level. We nonetheless analyzed the data of females only and largely obtained the same results as we obtained from the complete data set. A stronger focus on sex difference can be performed in future studies with several fold more mice than applied here.

A single experimental paradigm cannot measure all features of behavior. Nonetheless, the current study builds on our previous comparisons of S4^Disc1−/−^ and S4^Disc1+/+^ mice across multiple measurements of sleep where no differences were found, indicating that the *Disc1* deletion does not have a strong impact on these behaviors of S4 mice. Although this may also be true for other behavior paradigms, one can only be certain by performing the experiment. But since time and financial limitations prevent most laboratories from doing such tests, our current and previous studies with the *Disc1* deletion does not support the notion that it causes a strong phenotype and scientists performing experiments and reading the literature should not assume that it does.

A final limitation of our study is that the incomplete deletion of *Disc1* in S4 mice means that we cannot extrapolate what the effect of a complete disruption of *Disc1* would be.

In conclusion, this study characterized behavioral differences in S4 mouse strains presenting a natural *Disc1* mutation compared to the well-established B6 model in a seminatural environment. The *Disc1* deletion, previously reported to be involved in the expression of many molecular and physiological proteins, did not contribute to the largely different social behavior phenotypes studied here.

## Acknowledgments

This work was supported by the Knut and Alice Wallenberg Foundation, project no. 2019.0047, a fellowship from the Wallenberg Center for Molecular Medicine, and Konung Gustaf V:s och Drottning Victorias Stiftelse.

