## Supplementary data for "The *Disc1* deletion common to many inbred mouse strains has negligible effects on social behavior of 129S4 mice in a semi-natural environment"

### Supplementary information

### Supplementary figure 1

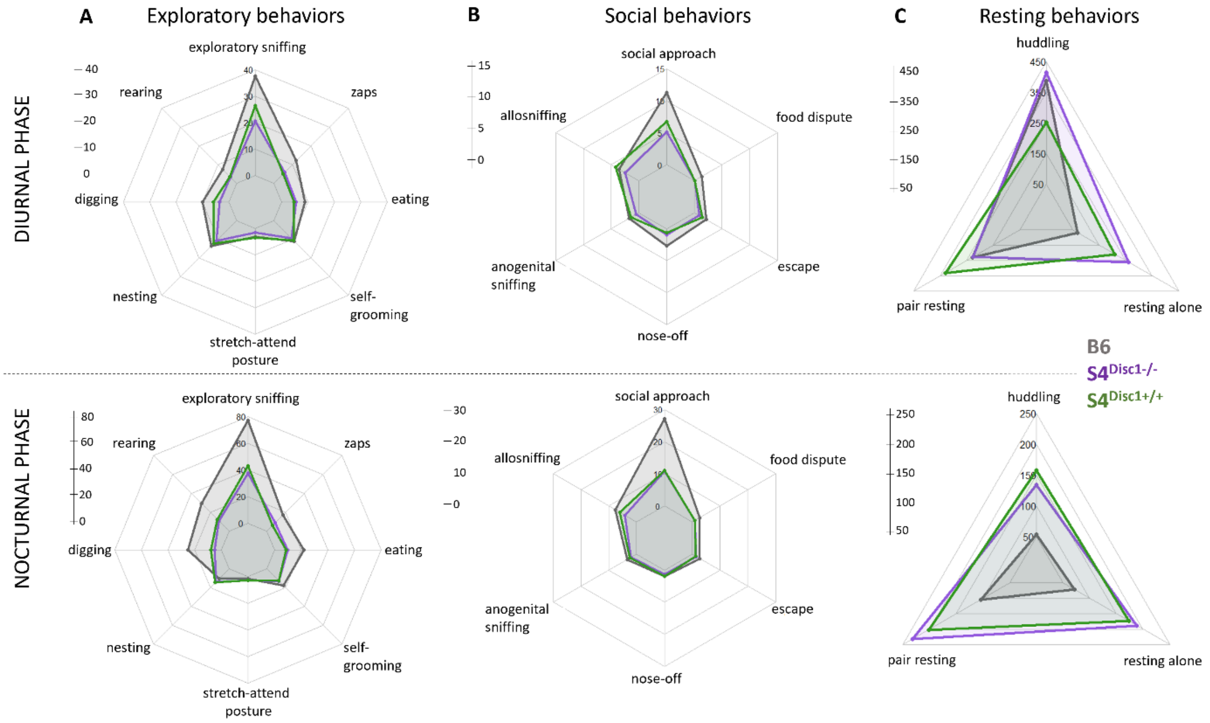


**Figure S1.** Behavioral profiles of **B6** (dark grey), **S4**$^{Disc1-/-}$ (**purple**) and **S4**$^{Disc1+/+}$ (**green**) mice during the diurnal and nocturnal phases of the light cycle. A. Average number of occurrences of exploratory and maintenance behaviors. B. Average duration of resting behaviors (s). C. Average number of occurrences of social behaviors.

**Supplementary figure 2**

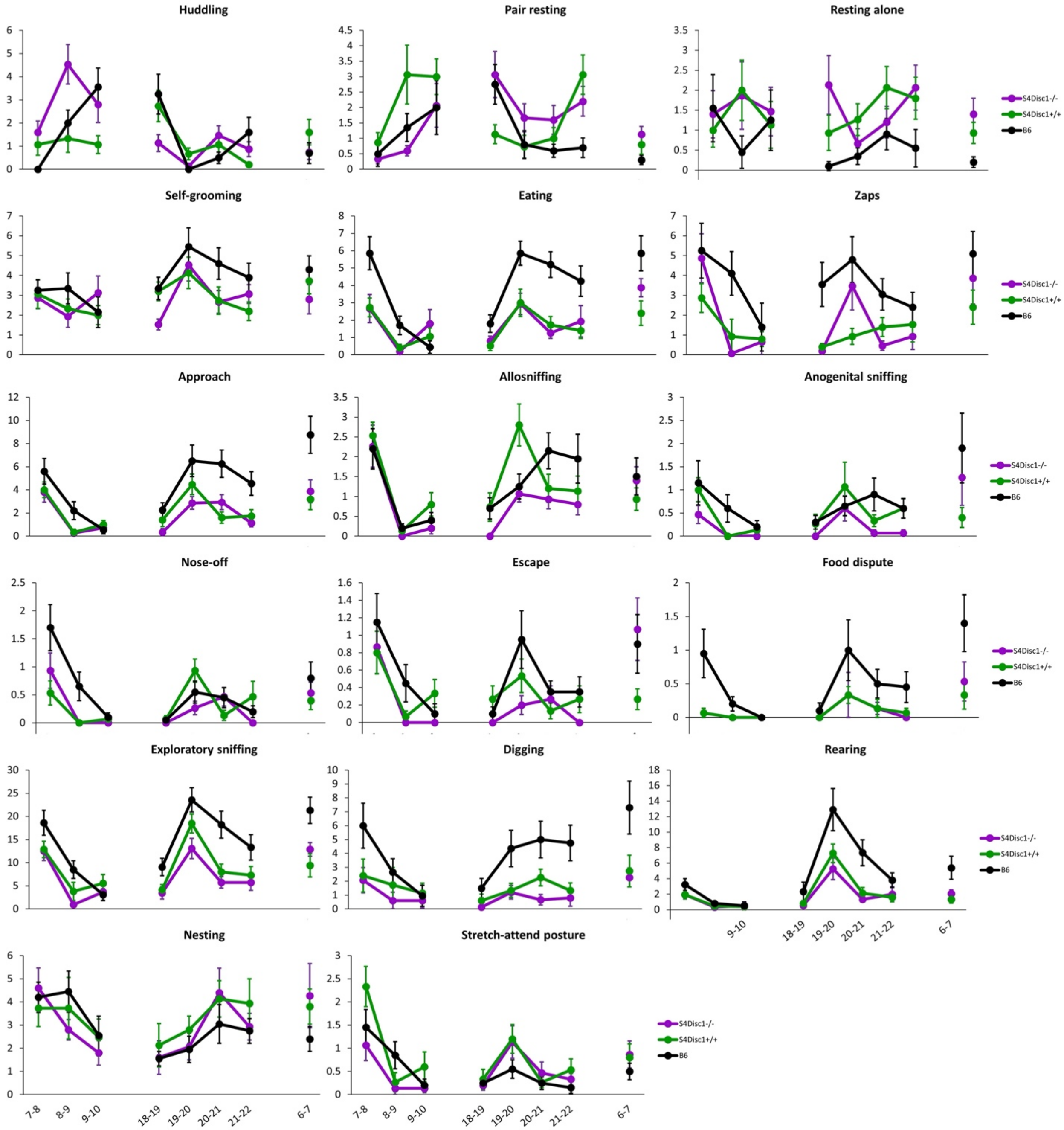


**Figure S2.** Behavioral occurrences per hour, over the 8 hours of observation. Room lights were on from 7:00 to 19:00.

**Supplementary figure 3**

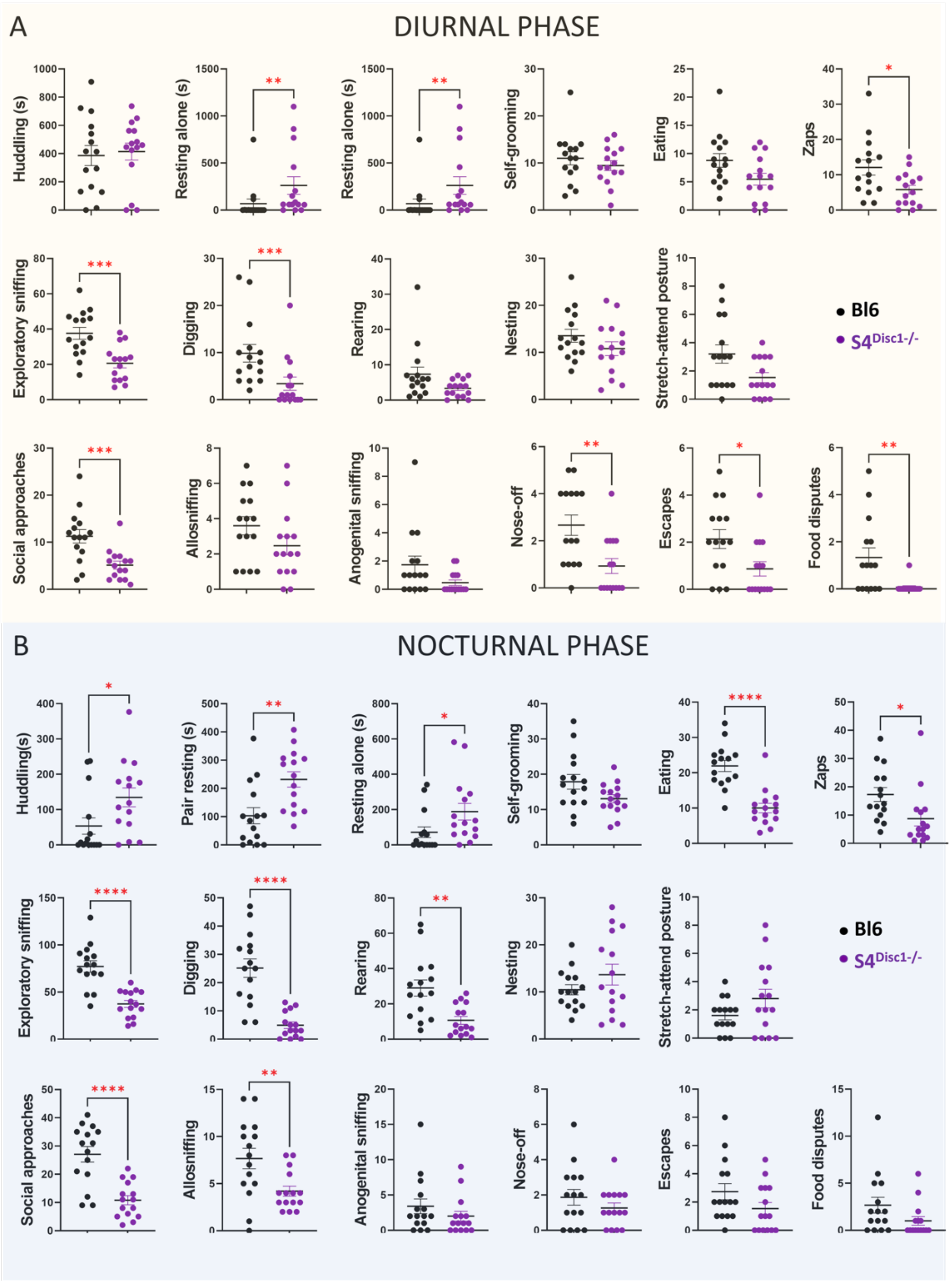


**Figure S3.** Comparison of maintenance, exploratory and social behaviors in natural S4$^{Disc1-/-}$ mutants (n=15) and B6 mice (n=15). Data are behavioral occurrences, except for resting behaviors (huddling, pair resting and resting alone), expressed in duration (s). Data are mean ± SEM. Student's *t*-test and Mann-Whitney test, * $p < 0.05$, ** $p < 0.01$, *** $p < 0.001$, **** $p < 0.0001$.

**Supplementary figure 4**

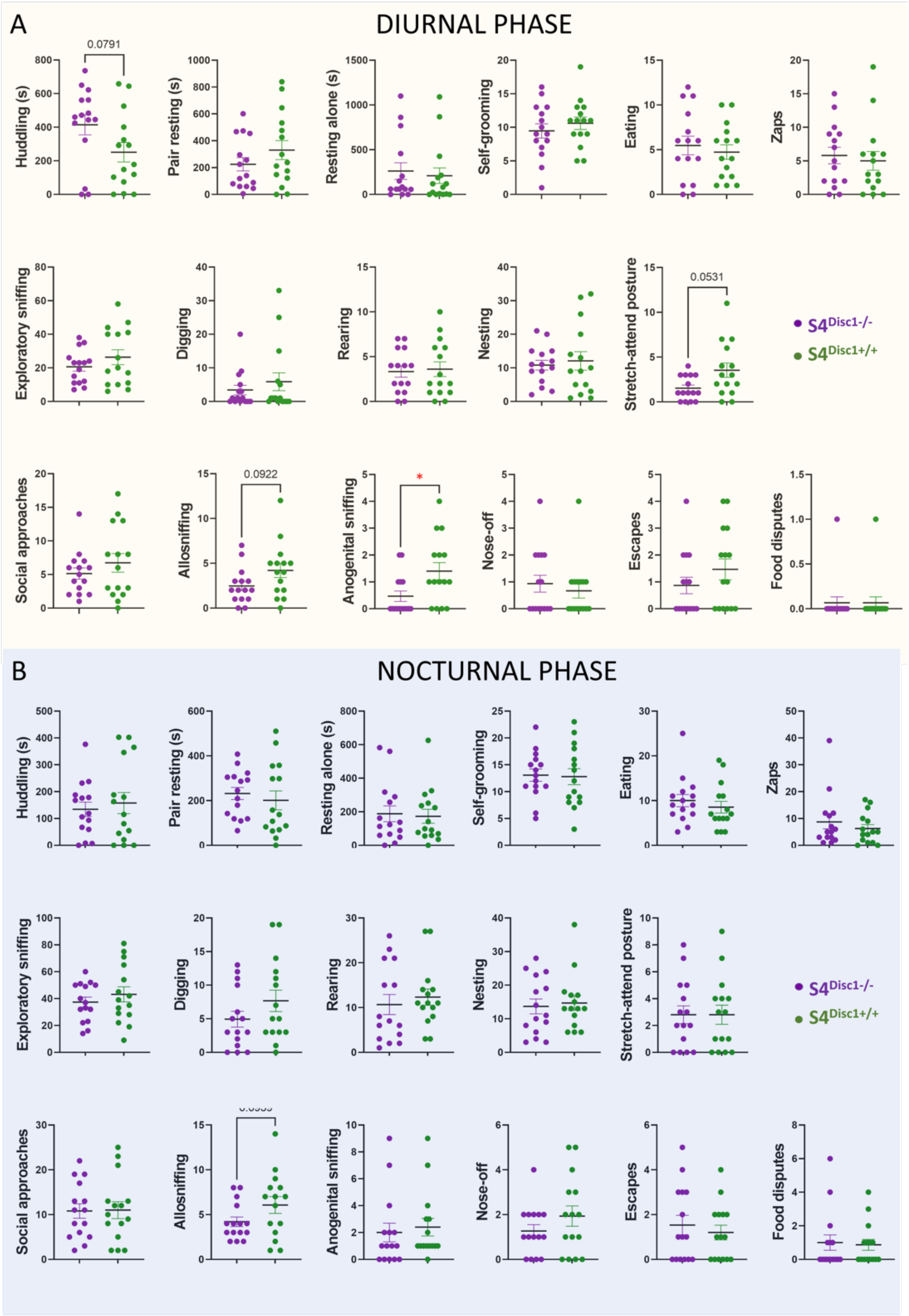


**Figure S4.** Comparison of maintenance, exploratory and social behaviors in natural S4$^{Disc1-/-}$ mutants (n=15) and S4$^{Disc1+/+}$ mice (n=15). Data are behavioral occurrences, except for resting behaviors (huddling, pair resting and resting alone), expressed in duration (s). Data are mean ± SEM. Student's *t* test and Mann-Whitney test, * $p < 0.05$. p-values inferior to 0.10 are indicated on plots.

Table S1. A summary of results presented in Figure 2 which includes data from males and then reanalyzed after data from males were excluded

| Behavior | Males included | | | Males excluded | | |
|---|---|---|---|---|---|---|
| **Diurnal** | $S4^{-/-}$ x B6 | $S4^{-/-}$ x $S4^{+/+}$ | $S4^{+/+}$ x B6 | $S4^{-/-}$ x B6 | $S4^{-/-}$ x $S4^{+/+}$ | $S4^{+/+}$ x B6 |
| Huddling | | | | | * | |
| Resting alone | * | | | | | |
| Eating | | | * | * | | * |
| Zaps | * | | | * | | ** |
| Exploratory sniffing | ** | | | * | | |
| Digging | ** | | * | ** | | |
| Social approaches | ** | | * | * | | |
| Nose-off | * | | ** | * | | * |
| Food disputes | ** | | ** | ** | | ** |
| Escapes | | | | ** | | |
| **Nocturnal** | | | | | | |
| Huddling | | | | * | | * |
| Pair resting | ** | | | | | |
| Resting alone | * | | * | * | | * |
| Eating | **** | | **** | **** | | **** |
| Zaps | * | | ** | | | ** |
| Exploratory sniffing | **** | | *** | *** | | *** |
| Digging | **** | | ** | **** | | **** |
| Rearing | *** | | ** | ** | | ** |
| Social approaches | **** | | **** | **** | | **** |
| Allosniffing | * | | | | | |
| Escapes | | | | * | | |

Data are mean ± SEM. One-way ANOVAs or Kruskal-Wallis tests, * $p < 0.05$, ** $p < 0.01$, *** $p < 0.001$, **** $p < 0.0001$
